# The secreted metabolome of HeLa cells under effect of Crotamine, a cell-penetrating peptide from a rattlesnake using NMR-based metabolomics analyses

**DOI:** 10.1101/2022.01.15.476435

**Authors:** Monika Aparecida Coronado, Fábio Rogério de Moraes, Bruna Stuqui, Marília Freitas Calmon, Raphael Josef Eberle, Paula Rahal, Raghuvir Krishnaswamy Arni

## Abstract

Sequestering and reprogramming of cellular metabolism represents one of the principal hallmarks several cells. Antimicrobial peptides have been shown to exhibit selective anticancer activities. In this study, the secreted metabolome of HeLa cells under action of the antimicrobial peptide Crotamine was evaluated. Although, Crotamine has been shown to be selective for highly proliferating cells and is able to extend the *in vivo* lifespan. The present study using a cell line of cervical cancer, HeLa cells provide insights into how Crotamine acts in cell metabolism. NMR spectroscopy was used to identify and quantify relative metabolite levels, which are associated with Crotamine uptake. Statistical analysis reveals that Crotamine dramatically affects metabolites related to glycolysis, metabolism and biosynthesis of amino acids and pyruvate metabolism. The developed machine learning model is found to be robust by ROC curve analysis, suggesting that the metabolic state of HeLa cells treated with Crotamine is different from the control samples. To account for metabolite levels, it is suggested that Crotamine would have to act on glycolysis, which, in turn, affects several other metabolic pathways, such as, glutathione metabolism, TCA cycle and pyruvate metabolism. The observed metabolic changes shed light into the mode of Crotamine function.

## 1. Introduction

Diverse factors such as exposure to carcinogens, viruses, ionizing radiation, chemicals as well as genetic disorders such as cell line mutations trigger the onset of cancer [1]. The Global Burden of Disease (GBD) estimated in 2015, there were 17.5 million cancer cases worldwide and for 2018 it was estimated to result in 9.6 million deaths. The statistics indicate that the incidence of cancer increased about 33% between 2005 and 2015 [2].

A non-enveloped small (8000bp) DNA-virus with circular double-stranded DNA, has been implicated in causing genital warts and cancer. More than 40 years ago, Human Papillomavirus (HPV) was implicated in causing different human neoplastic lesions [3] and currently, 200 HPV types including cutaneous and mucosal HPV types have been characterized. Cervical cancer (CC) is certainly the most common cancer among women causing significant morbidity and mortality worldwide [4–6]. In 2012, it was estimated that more than 527,000 new cases with an 85% occurrence rate was registered in less developed regions resulting in and more than 265,000 deaths with 87% of them from less developed countries [6, 7]. In 2018, it was estimated for almost 570,000 and the numbers of deaths was more than 311,000 worldwide with an estimation to increase more than 35% from 2018 to 2040 [8, 9]. At the global level, one in 68 women develops CC in their life-times, the occurrence is higher in developing countries, with 1 in 24 women presenting CC symptoms, and is significantly lower in highly developed countries, where 1 in 115 women developed CC during their lifetimes [2].

According to the National Cancer Institute of Brazil (INCA), the estimated number of new cases of CC for each year of the biennium 2018-2019 was estimated with a risk of 16,370 cases per 100.000 women in Brazil. It represents the seventh neoplasm more frequent in female population in the world ranking, and the fourth cause of women’s mortality by cancer in Brazil [10].

Proteins, peptides and enzymes from animals of different species are being tested for use in cancer therapies and many active secretions produced by animals have been employed in the development of new drugs to treat diseases such as hypertension and cancer. Snake venom toxins have contributed significantly to the treatment of many medical conditions [11] and a host of studies cite the anti-cancer potential of snake venoms [12,13].

Since treatment of cancer is a major challenge and many of the currently used therapies are prohibitively expensive and often trigger undesirable secondary reactions. Molecules isolated from snake venoms have been demonstrated to retard and inhibit the growth of cancerous cells and hence have been the focus of research [12]. Crotamine, extracted from the venom of the Brazilian rattlesnake, *Crotalus durissus terrificus*, is a highly basic (pI 10.3), low molecular weight (42 kDa), non-enzymatic and, non-cytolytic cell-penetrating peptide (CPP) [14–16] that can cross the cellular lipid barrier [17–19]. It also exhibits other activities such as: antimicrobial activity [20,21], selectivity for highly proliferating cells [20,22]; inhibits tumor growth [23], and it is also potentially interesting as an individual therapeutic agent since it possesses the ability to transport proteins, peptides, nucleic acids and, perhaps, even entire genes across the cellular membrane [17,24–28].

Selective cytotoxicity of Crotamine in tumor cells has been reported: experiments using Crotamine as an inhibitor resulted in a significant decrease of tumor growth and, mice with tumors showed an increase in survival rates [23,29] and it has also been demonstrated to possess anti-tumoral activity in cell cultures and animal models [22,23,29,30]. Experiments using *in vitro* model are more feasible and practical to assess the potential effect of molecules and to understand the underlying mechanisms of physiological processes. Understanding the cellular responses of Crotamine may provide metabolic information on the reactions of biological systems at the molecular level. The biological interactions in cellular pathways can be elucidated based by the analysis of cell lines using NMR-based metabolomics foot printing assays, to screen promising drugs, and is also effective in *in vitro* screening for identifying significant metabolite changes in response to the administration of diverse molecules [31–33].

In the present study, we utilized an integrated metabolomics approach to systematically investigate the response of Human cervical adenocarcinoma cells (HeLa cell line) using Crotamine as the mediator. Based on the analysis of the metabolic pathway, we identified changes of the cell line pattern. 1H NMR spectroscopy coupled with pattern recognition and biochemical network analysis was used to characterize the metabolic footprint profile of HeLa cells.

## 2. Materials and Methods

### 2.1 Cell Line

Human cervical adenocarcinoma cell line (HeLa) was kindly provided by Luisa Lina Villa (Department of Radiology, Center on Translational Oncology Investigation, São Paulo State Cancer Institute, São Paulo University, Brazil). Cell lines were cultured in Dulbecco’s modified Eagle medium (DMEM) (Gibco Life Technologies, Grand Island, NY, USA) supplemented with 10% heat inactivated fetal bovine serum (FBS), penicillin (50 U/mL), and streptomycin (0.05 mg/mL). Cells were incubated at 37 °C with 5% CO2.

### 2.2 MTT Assay

Colorimetric MTT assay was used to evaluate the cytotoxicity in 96-well plates and the medium containing Crotamine at concentrations of 6, 8, 10, 12, 14, 16, and 20 μM was added to each well containing 1 × 104 cells. After 4, 12, and 24 h of incubation with Crotamine, 100 μL of medium containing 3-(4,5-dimethylthiazol-2-yl)-2,5-diphenyl tetrazolium bromide (MTT) (1 mg/mL) (Sigma-Aldrich) was added to each well. After 30 min of incubation, the medium was removed, and the formazan crystals were solubilized by incubation for 10 min in 100 μL of DMSO (Sigma-Aldrich). Absorbance of each well was determined at 570 nm. Each experiment was performed in triplicate and in three independent assays.

### 2.3 Crotamine treatment and Metabolite Extraction

For cell seeding, HeLa cells were cultured in serum-starved medium. After 24 h, Crotamine was added to the supplemented culture medium (containing 10% FBS and antibiotics) at concentrations of 10 μM. Crotamine treatment concentration was chosen based on the results of the MTT assay. To avoid contamination in the cell culture, filter sterilization (using a Millipore filter with a pore size of 0.22 μm) of the medium containing Crotamine was applied before introducing the medium to the cell culture. After 24 hours of Crotamine incubation, the medium was submitted to footprint metabolomics studies. The control group received only serum starved medium by 24 h.

### 2.4 Nuclear Magnetic Resonance Spectroscopy

Nuclear Magnetic Resonance (NMR) measurements from sample cultures were performed in a Bruker AVANCE III HD (Germany), operating at 600 MHz for 1H equipped with a triple resonance cryoprobe. The standard NOESYPR1D pulse sequence was used with a recycle delay of 2 s, a mixing time of 100 ms, 16 scans (four dummy scans), collected in 32 k complex data points and a spectral width of 20 ppm. Each free induction decay measurement (FIDs) was multiplied by an exponential function with 1 Hz line-broadening, followed by Fourier transformation in the Bruker TopSpin 3.2.5 (Bruker Biospin, Germany). Reference in each spectrum was performed internally by setting the lactate methyl doublet at 1.31 ppm. Finally, the residual water region (4.5-5.0 ppm) was deleted from the dataset.

Furthermore, specific regions were formerly subjected to integral calculation, after metabolite identification, for relative concentration over samples. For assisting metabolite identification, selected samples were used for collecting 1H, 13C Heteronuclear Single Quantum Coherence spectroscopy (HSQC).

### 2.5 Statistical Analysis

Data was organized following MetaboAnalyst guidelines [34] and its web server was used for analysis. First, data was Pareto scaled (mean centered and divided by the square root of the standard deviation), and subsequently, Principal Component Analysis (PCA), Partial Least Square Discriminant Analysis (PLS-DA) and its orthogonal version (OPLS-DA) were performed. First, the complete spectra (excluding the solvent region) were uploaded for evaluating control and treated metabolic differences by means of PCA, PLS-DA and OPLS-DA score plots. Significant signals related to high absolute loadings were subject to metabolite identification. Specific metabolite signals quantified by their integrals were reorganized and uploaded to MetaboAnalyst for further analysis. Univariate t-test and fold change analysis were used to compare relative metabolite levels between control and Crotamine treated HeLa cells. Boxplots and t-test p-values were used to check differences in metabolite levels. Also, PCA, PLS-DA and OPLS-DA were repeated with this reduced dataset to confirm if the used metabolites were able to keep the degree of difference between control and Crotamine-treated cell lines. Furthermore, machine learning classification methods, Random Forest and Support Vector Machine (SVM) were used to check if the developed models were successful in predicting whether cells were from a control or Crotamine-treated sample.

To assess the robustness of the developed model with respect to metabolite levels following Crotamine treatment, Receiver Operating Characteristic (ROC) curves for three different classification algorithms, as available in MetaboAnalyst at Biomarker Analysis module was used. For each model, the web-server uses a Monte-Carlo cross validation scheme where 66% of the data is used to assess feature importance and build the model itself, whereas the remaining 33% are used for validation and performance measurement. This process is then repeated using different metabolites sets. Metabolite importance is assessed according to the frequency of occurrence of a particular metabolite is selected to compose a classification model.

### 2.6 Metabolite Identification

Metabolites were manually identified using Chenomx profiler (version 8.1, Chenomx Inc., Canada) by matching resonances of random spectra. Cross-peaks were identified in the TOCSY spectra as well as signals from the 1H, 13C-HSQC. Expected TOCSY and HSQC signals were retrieved from the Human Metabolome Data Bank [35].

### 2.7 Pathway Analysis

Genome-scale metabolic models have long been used for gaining insights into the response of metabolic pathways of a variety of different stimuli in context to an organism, a key concept in the system biology approach [36]. HeLa cell lines metabolic model was generated by using RNA-seq data and evaluated with metabolome mapping, showing interesting results for the identification of cancer related metabolites in different cell lines [37]. In this study, HeLa cells metabolic model were uploaded into MetExplore web server [38] and the observed metabolite levels were used to get insights into the metabolic path-ways that were changed by the uptake of Crotamine.

## 3. Results

### 3.1. Viability of HeLa cells in response to the exposure of Crotamine

Crotamine was purified following the procedure of Coronado et al. (2013) [16] and used for cell viability assays. Experimental control cells retained their viability at all analyzed incubation times. After 4, 12 and 24 h of incubation with all the tested concentrations of Crotamine, HeLa cells demonstrated viabilities greater than 70% (Figure 1). Viability of HeLa cells decreased with increasing Crotamine concentration; however, the concentrations used in this work demonstrated low cytotoxicity. From these results, the 10 μM concentration and 24 h incubation were chosen, as the results showed great cell viability of around 90%.

**Figure 1.**
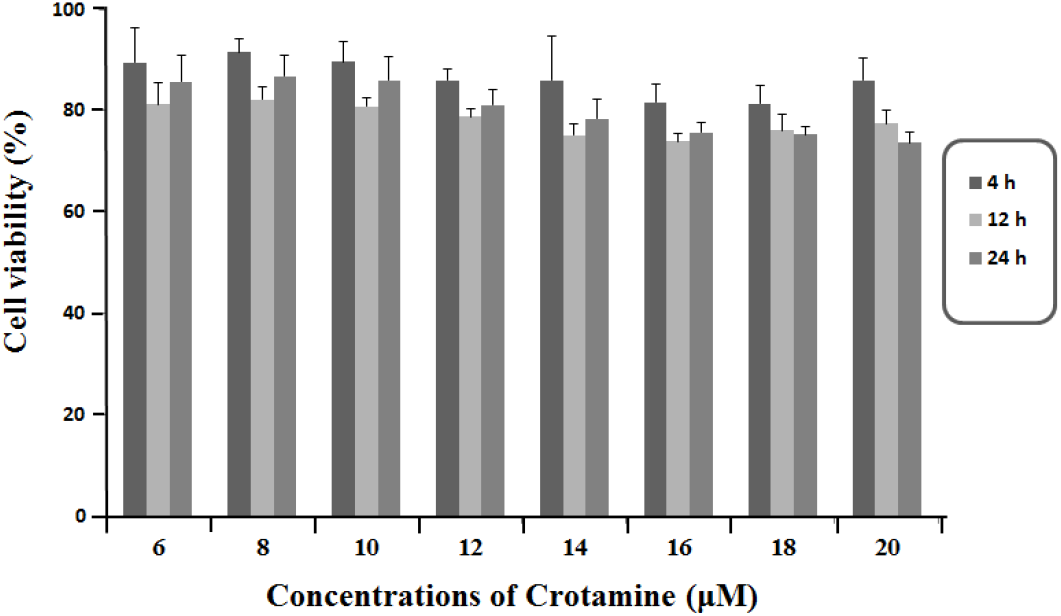
Analysis of the cellular effects of Crotamine on HeLa cells viability. Cell viability assay investigated after 4, 12 and 24 h of incubation with Crotamine at concentrations of 6, 8, 10, 12, 14, 16, and 20 μM.

### 3.2. Secreted metabolic profile of HeLa cells

Since the highest cell viability obtained on HeLa cells was after 24 h, we decide to evaluate the secreted metabolomic profiling on HeLa cells treated with 10 μM Crotamine and to compare this with the results for untreated cells.

For the present study, metabolic foot printing was been used to characterize the metabolic changes of HeLa cells subjected to Crotamine. Intensities from NMR spectra were examined, and spectral bins were manually selected to exclude noisy and solvent regions.

Following the treatment of the Crotamine NMR profile of HeLa cells, the observed metabolic differences between control and treated cells (Figure 2-asterisk), indicated that Crotamine induced alterations in the metabolism of HeLa cells.

**Figure 2.**
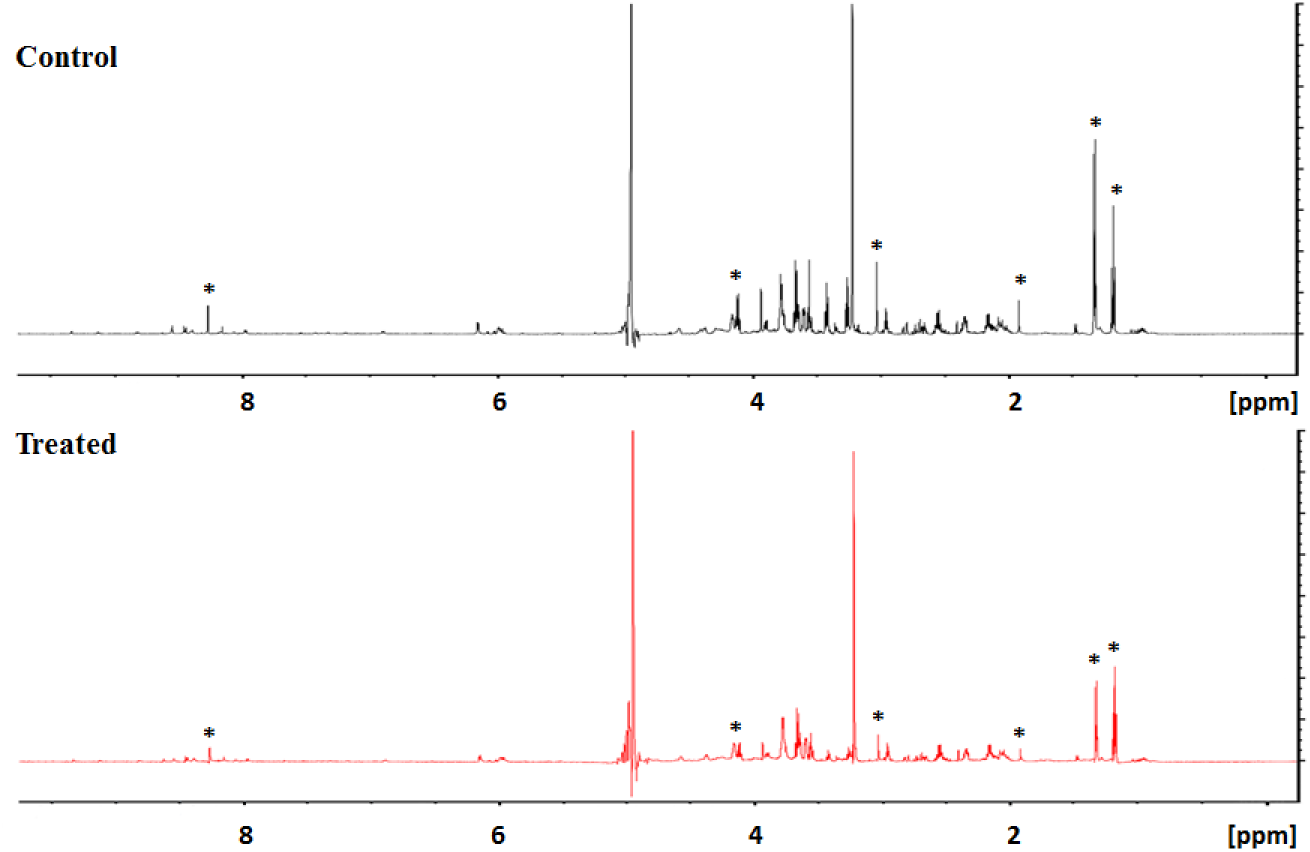
NMR spectrum comparison. NMR spectrum comparison between control and treated sample.

The analysis of the spectra shows the presence of signals (i) leucine, isoleucine, valine, ethanol, lactate, pyruvate, lysine, pyroglutamate, threonine, and glucose in the spectra region from 0.0 ppm to 5.5 ppm, (ii) 1-methylhistidine, tyrosine, phenylalanine, and formate in the spectra region from 7 ppm to 8.5 ppm (Table1; Figure 3).

**Figure 3.**
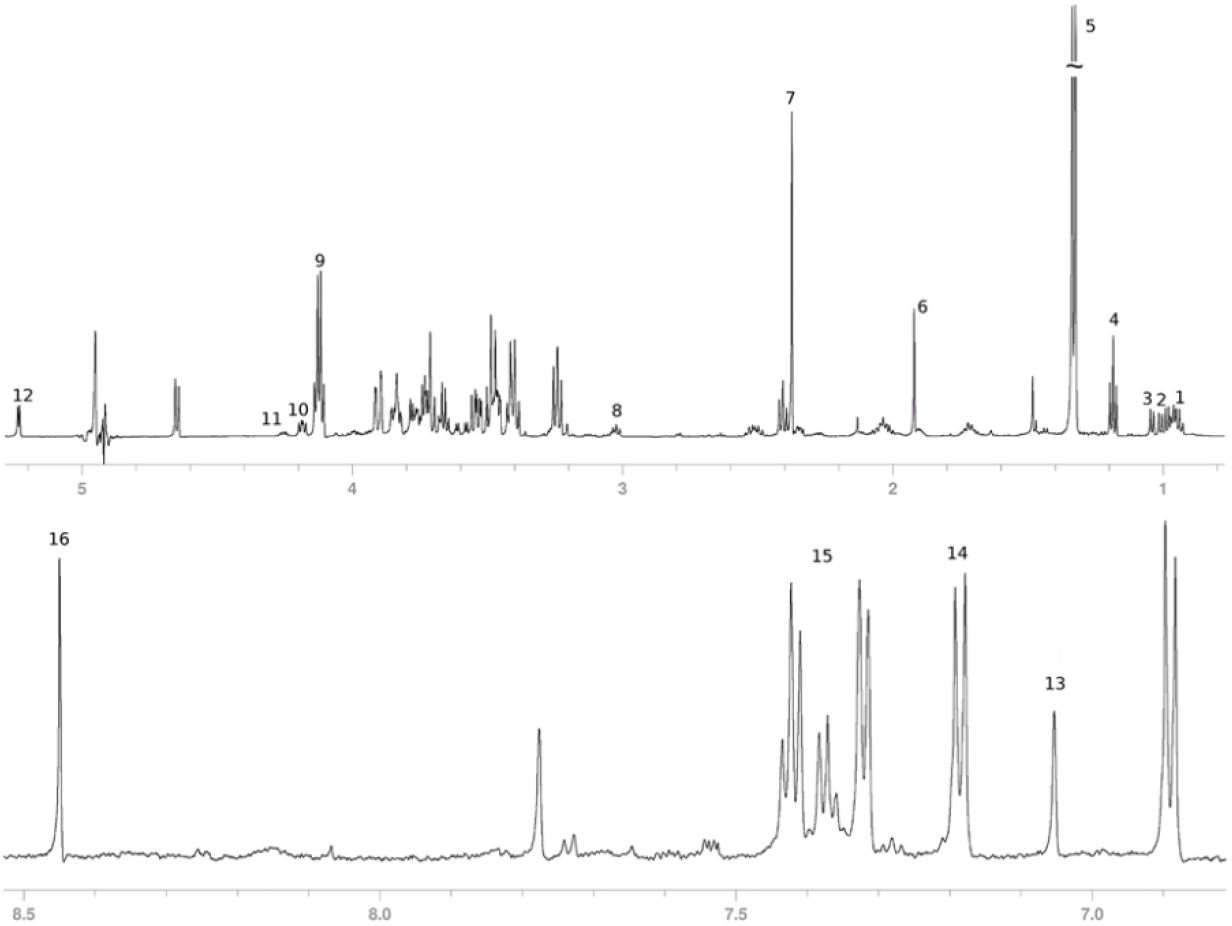
Sample NMR spectrum. NMR spectrum showing signals for: 1 – Leucine; 2 -Isoleucine; 3 – Valine; 4 – Ethanol; 5 and 9 – Lactate; 6 – Acetate; 7 – Pyruvate; 8 – Lysine; 10 – Pyruglutamate; 11 – Threonine; 12 – Glucose; 13 – 1-Methylhistidine; 14 – Tyrosine; 15 – Phenylalanine; 16 – Formate.

**Table 1.** List of HeLa metabolic pathways affected by Crotamine uptake, using a p-value of Fisher’s test >5%.

| Pathway | Coverage | # Metabolites matched | Right tailed Fisher-test |
| --- | --- | --- | --- |
| Transport of extracellular metabolites | 2.13 | 13 | 7.76E-11 |
| Protein degradation | 12.73 | 7 | 1.04E-10 |
| Aminoacyl-tRNA biosynthesis | 11.11 | 7 | 2.81E-10 |
| Artificial reactions | 8.97 | 7 | 1.31E-09 |
| Transport to mitochondria | 3.56 | 9 | 9.39E-09 |
| Transport lysosomal | 5.26 | 7 | 5.71E-08 |
| Glycolysis/Gluconeogenesis | 7.81 | 5 | 1.01E-06 |
| Transport peroxisomal | 3.88 | 4 | 2.38E-04 |
| Pyruvate metabolism | 6.12 | 3 | 4.35E-04 |
| Valine, Leucine and Isoleucine metabolism | 4.76 | 3 | 9.13E-04 |
| Glutathione metabolism | 5.56 | 2 | 5.58E-03 |
| Tyrosine metabolism | 5.56 | 2 | 5.58E-03 |
| Phenylalanine, tyrosine and tryptophan biosynthesis | 2.5 | 3 | 5.80E-03 |
| Transport Golgi apparatus | 3.64 | 2 | 1.27E-02 |
| Cysteine and methionine metabolism | 3.03 | 2 | 1.80E-02 |
| Glycine, serine and threonine metabolism | 2.3 | 2 | 3.02E-02 |

### 2.3. Crotamine induced metabolic variation in HeLa cells. (Multivariate data analysis)

Multivariate data analysis was employed to analyze the changes caused by Crotamine on HeLa cells and to identify the possible metabolic pathway involved.

PCA, PLS-DA and OPLS-DA score plots show well selected controls and the treated groups in the 95% confidence interval, indicate significant metabolic changes in HeLa cells because of the Crotamine treatment. Principal component analysis (PCA) of the 1H NMR spectra showed a clear discrimination between HeLa cell line before and after 10 μM treatment with Crotamine (Figure 4a). To obtain information on the types of metabolites responsible for the class separation, the orthogonal projection to latent structure with discriminant analysis (OPLS-DA) was conducted with the corresponding NMR data from the cell group. As illustrated in Figure 4b, the control and Crotamine-treated groups could be clearly distinguished in the OPLS-DA score plot.

**Figure 4.**
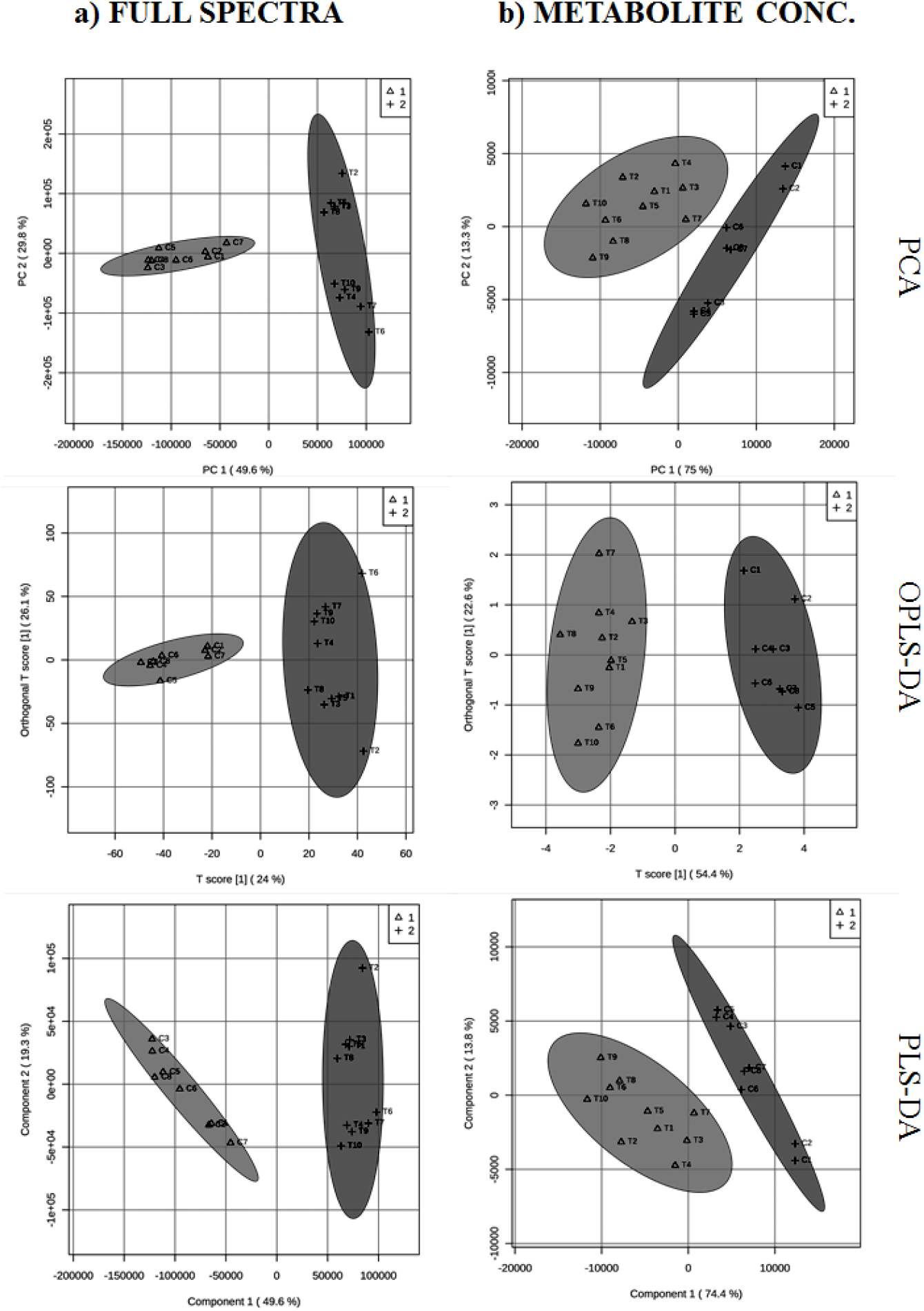
PCA, PLS-DA and OPLS-DA score plots. Principal component analysis (a), orthogonal partial least square discriminant analysis (b) and partial least square discriminant analysis score plots performed for full spectra bins and integral signals for selected metabolites.

Additionally, signals that satisfy the requirements of possessing a Welch-test p-value lower than 5%, a fold change in log scale higher than 1.2 or lower than 0.8 and have high VIP score from PLS-DA loadings were selected (Figure 4c). These signals were used to metabolite identification as performed by matching chemical shifts and the scalar coupling in the Chenomx NMR software package and 1H,13C-HSQC signals. A total of 15 metabolites, presented in the Table 1 and in Figure 3, were considered relevant for distinguishing between control and Crotamine treated cell cultures. Their levels encountered in Crotamine-treated and control samples were measured using the integral module in TopSpin 3.2 for isolated signals of each metabolite (Fig. 4 Metabolite conc.). The measured values were scaled in MetaboAnalyst to identify which metabolic pathways are involved. The tool suggests the most relevant pathways by uploading the discriminatory compounds that were significantly influenced by Crotamine treatment. Results are displayed in figure 5 as boxplots.

**Figure 5.**
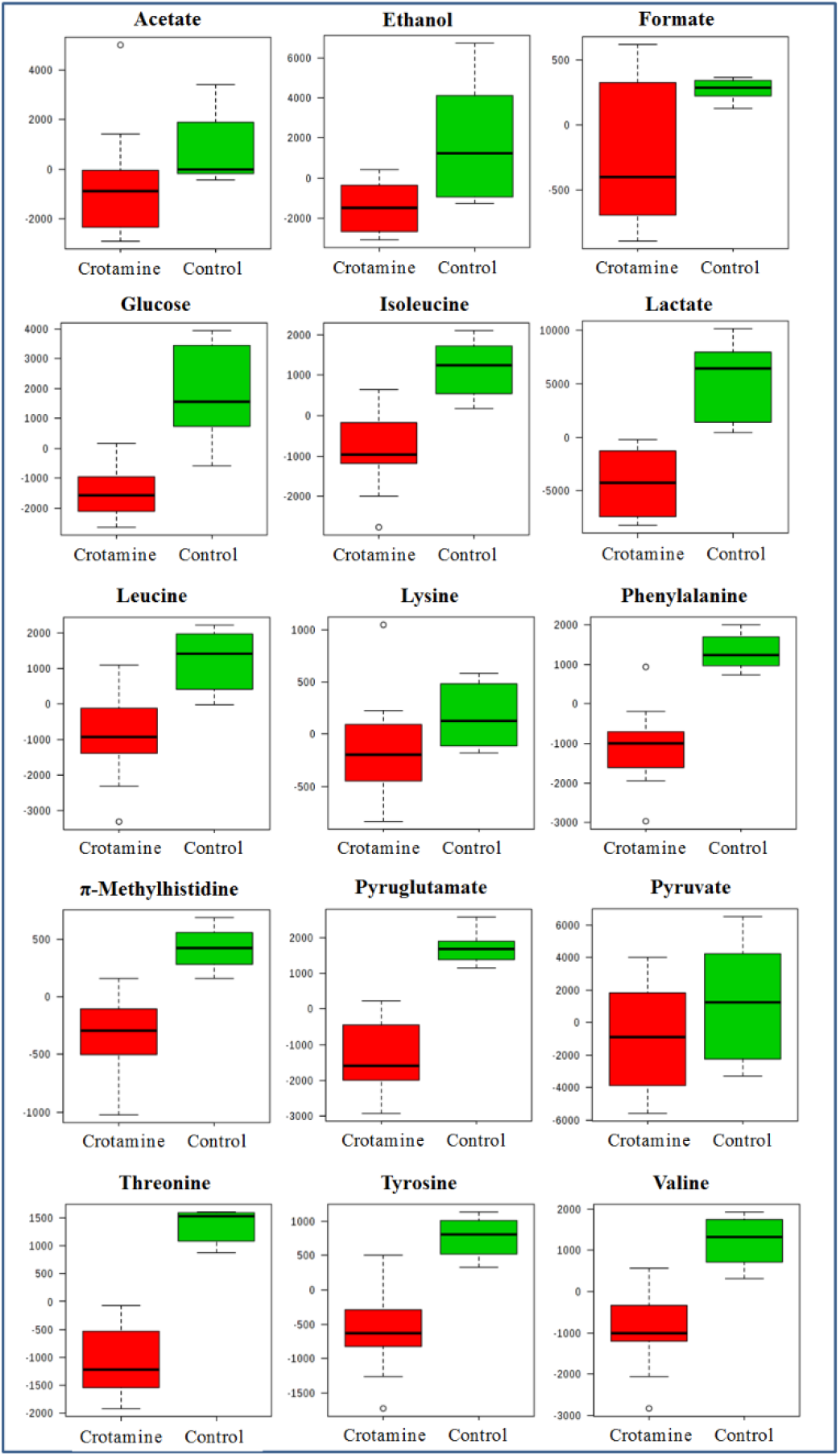
Metabolite levels in Crotamine-treated and control samples. Levels of each metabolite found to be important for distinguishing between Crotamine-treated and control samples. Values were evaluated from integrated isolated signals for each metabolite presented in NMR spectra and scaled in MetaboAnalyst.

In our study, the stress-related metabolic variations include the decreases of acetate, ethanol, formate, glucose, etc. The decreased levels of amino acids such as isoleucine, leucine, lysine, phenylalanine, threonine, tyrosine, and valine indicate an inhibitor of protein metabolism in response of Crotamine. Lactate and acetate release are closely correlated with the variance of the glucose utilization mainly due to cytosolic pyruvate production in glycolysis.

Each of the models developed to analyze the data provides an overview of the metabolite enrichment set as presented in figure 6a and ranked according to metabolite importance. Classification probabilities for each sample following the Monte-Carlo cross validation is shown in figure 6b. Perfect classification is achieved for both SVM (3 metabolite levels used) and PLS-DA (2 metabolite levels used). These results indicate that the metabolite levels used in the models can potentially applied to differentiate between the metabolic states of control HeLa cells and following Crotamine treatment.

**Figure 6.**
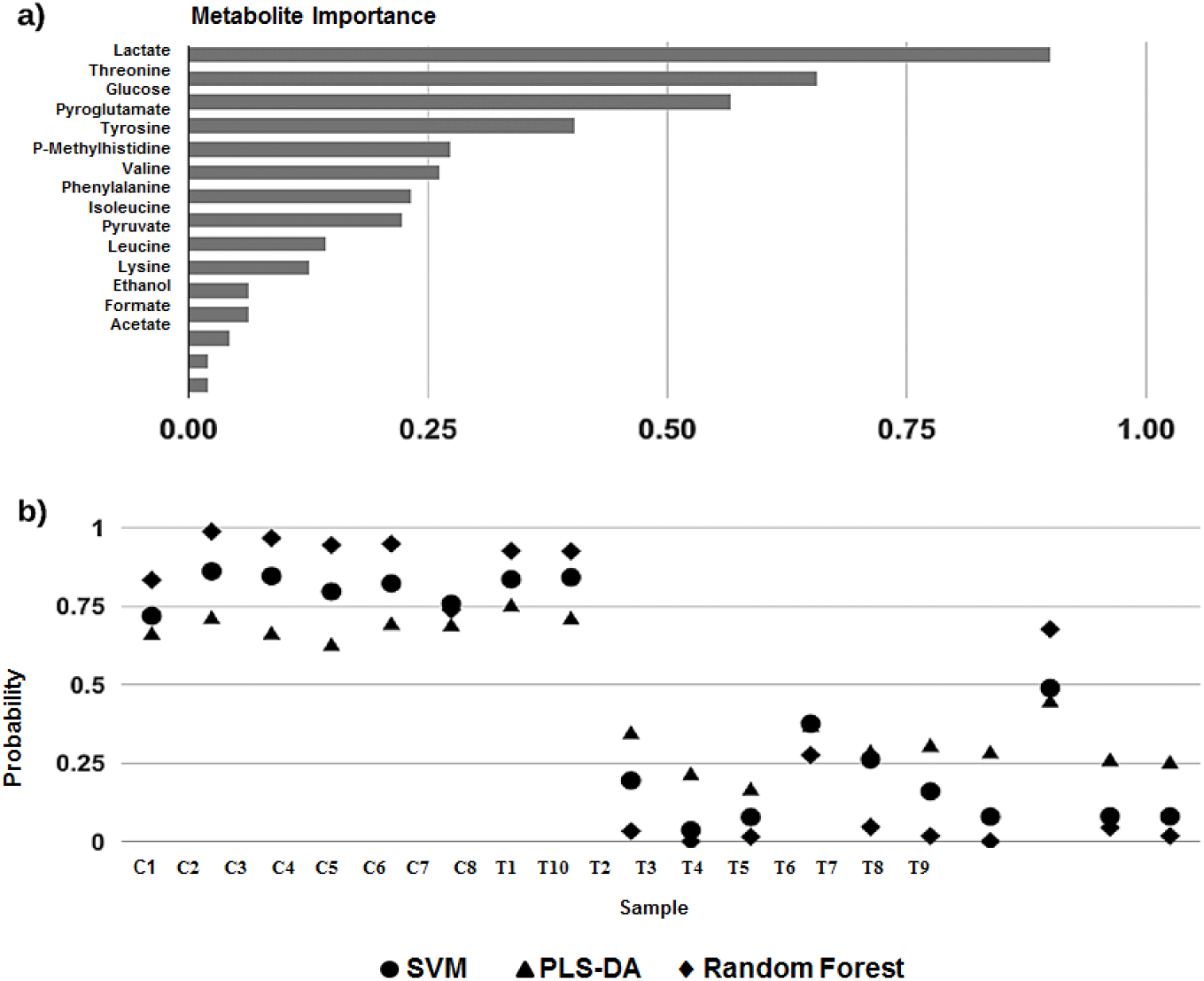
Metabolite Importance and Classification Models. Metabolite importance measured as the average of used frequency in classification models SVM, PLS-DA and Random Forest (a). Classification probabilities for each sample tested in Monte-Carlo cross validation by classification models SVM, PLS-DA and Random Forest (b).

## 4. Discussion

In order to identify the pathway and to identify the roles in cancer development, the observed metabolite levels were analyzed using genome scale metabolic models for HeLa cells that were uploaded into the MetExplore web server and based on the metabolome analysis module, the metabolite levels measured by NMR were set and a list of affected pathways were retrieved using a right tailed Fisher’s test (Table1). Since some metabolite levels were reduced in the Crotamine treated samples, transport reactions can be recovered to be altered from non-essential metabolites and between different compartments within cell. Due to the observed changes in the amino acids levels, protein degradation and aminoacyl-tRNA biosynthesis pathways are evaluated by MetExplore analysis. Artificial reactions are a set of chemical reactions and account for biomass content (amino acids, lipids and nucleotides), vitamins derivatives and cofactors available in the model and, for instance, are not of special importance. Interestingly, the metabolic pathways found to be affected in HeLa cell treated with Crotamine are all interlinked as observed (Fig. 7).

**Figure 7.**
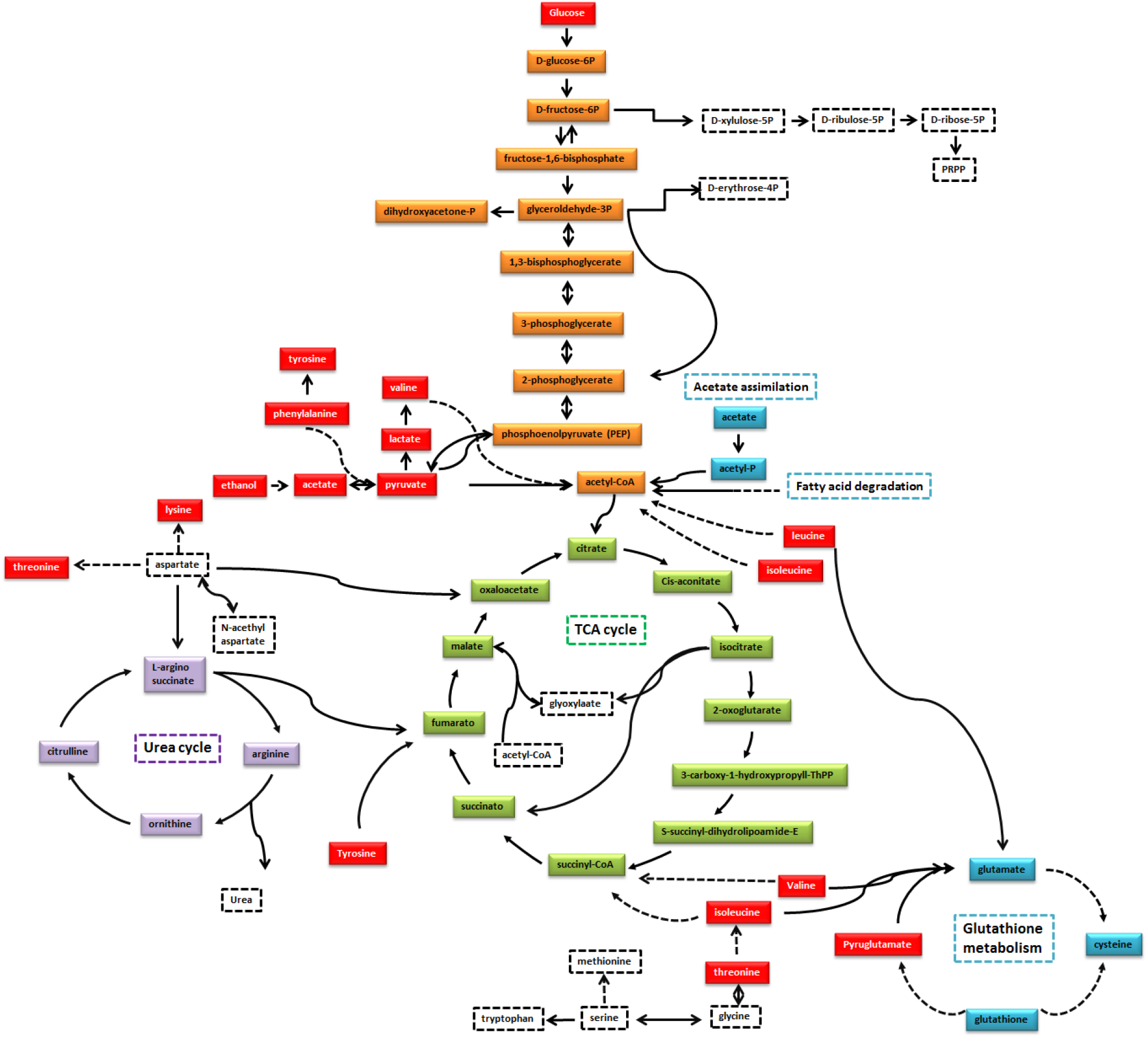
Important pathways affected by Crotamine in HeLa cells. Crotamine-treated samples were found to have altered levels of the metabolites in red. Important pathways retrieved by genome-scale metabolic modeling are shown. Straight lines show direct biochemical reactions while dashed lines represent biochemical steps not shown.

As mentioned earlier, many metabolites were affected by treatment with Crotamine. Furthermore, lactate is found to be the most used metabolite of the classification (Figure 6a). Independently of oxygen, most cancer cells produce their energy by taking up glucose and producing lactate, it was observed by Otto Warburg in the 1920s, in Warburgs effect [39–42], which increase the levels of lactate. The reduced lactate levels in Crotamine-treated samples are in line with drug effects that target glycolysis [43]. The same point holds for pyruvate levels, since one is the precursor to the other, and it is the end-point in glycolysis (Figure 7). Many studies indicate that tumors cells present the capacity to produce energy in the aerobic oxidation of glucose [40,44]. Furthermore, mitochondrial metabolism is fundamental in the process of cancer cell proliferation [45]. The TCA cycle can be supplied by other sources using fatty acid and amino acids as substrate to maintain ATP production in cancer cells [40,46].

Reduction in glucose levels, the third metabolite in the rank of importance, is observed in treated samples. Increase in glucose uptake may be achieved by overexpressing a set of proteins known as glucose transporters [47–49], which indicate that cells are changing their metabolic state in order to increase energy production that is affected by Crotamine. Glucose can be switched on by acetate, to support the biosynthesis of fatty acids and lipids, as demonstrated in tumor-like tissue culture condition [44,50,51]. One rescued acetate molecule can produce acetyl-CoA to synthesize fatty acid or sterols, to acetylation of histones, or to promote oxidation to generate additionally ~12 units of ATP, all of this process at the cost of a single ATP [52]. Some studies address the importance of acetate in cancer development [53–56]. One alternative source of carbon can be provided by acetate, as well as by glutamine if the access to the glucose-derived from acetyl-CoA is compromised by hypoxia or mitochondrial dysfunction [44, 56].

Dependent of an acetyl-CoA synthetase2 (ACSS2), studies using metabolomics demonstrated that cancer-cell can capture acetate molecules to use as a carbon source [53,57]. An acetate, when ligated to CoA is the most central and dynamic metabolite in intermediary metabolism (Fig.7).

It is well documented that cancer cells generate energy differently than normal cells [53,58,59]. The process that healthy cells get energy is very complex and requires oxygen molecules; however, the current knowledge indicates that this does not occur in cancer cells [44,60,61]. The metabolic pathway used for cancer cells to produce energy was demonstrated to be more primitive [44,47,60–61]. The less efficient process, the Warburg effect has been studied extensively. In the 1920s, several experiments performed by Otto Warburg demonstrated that cancer cells generate energy via fermentation, similar to the process observed in yeast, even when there is a sufficient amount of oxygen available to break down glucose [39, 62]. Using fermentation, which is less efficient, the cancer cells should use more glucose to produce the same amount of energy, suggesting that malignant cells have to rely on other sources for metabolism, like fats and amino acids [63]. Tyrosine is one of the sources that cancer prefers but is rarely used by normal healthy cells. Some *in vitro* studies demonstrate that the restriction in the availability of tyrosine, as also methionine, and phenylalanine affects the signaling pathway, and induces apoptosis in some cancer cells lines [64–66].

Many studies have shown that the restriction of specific amino acids modulates glucose consumption and the changes are closely related to glucose metabolism. In prostate cancer cells the carbohydrate metabolism is modulated by the restriction of amino acid, as shown by Fu et al., which causes modification in glucose metabolism, thereby leading to cell death and apoptosis [64,67].

Phenylalanine, one of the reduced molecules in our study is an essential amino acid and its hydroxylation produces tyrosine. The ketogenic component, which in fact, is one important component of the tricarboxylic acid (TCA) cycle, is produced by the degradation of tyrosine to acetoacetate and fumarate (Fig.7). Oxaloacetate, which is converted from the fumarate, can be directed to the gluconeogenesis pathway. There are five amino acids entering the TCA cycle via pyruvate, these are: alanine, glycine, serine, cysteine, and tryptophan. In Crotamine-treated samples we did not observed any modification in these amino acids. However, the other amino acids that yield acetyl-CoA and/or acetoacetyl-CoA, like as lysine, phenylalanine, tyrosine, leucine, and isoleucine are affected by Crotamine. On the other hand, threonine is one amino acid that can be converted in glycine and play a pivotal role in one-carbon metabolism and nucleotide synthesis [60]. However, this is an amino acid that cannot be synthesized by humans; all demand is fulfilled by uptake. Mammalian cells require threonine as an essential amino acid [68]. Threonine plays a key role in cancer cell growth and proliferation [69]. Cell death and reduction in methylation of some histone are results from the shortage of threonine in cell cultures [60,68]. There is a sequence of simultaneous events; tyrosine reduction, probably reduce the glutamate synthesis, which automatically reduce its PTM (post-translational modification), it means, the reduction of pyroglutamate, the cyclic form of glutamate.

Following the described metabolite changes, the cyclic amino acid pyroglutamate is encountered at the N-terminus of some protein and biological peptides [70], which is often involved in stabilizing the protein by making it more resistant to chemical or enzymatic degradation [71,72]. Pyroglutamate is an intermediate in glutathione metabolism, whereas, glutathione (GSH) plays an important role in a multitude of cellular processes. In cancer cells it is particularly relevant in the progression and regulation of carcinogenic mechanism. GSH is crucial to control the reactive oxygen and, its deficiency leads to a susceptibility to oxidative stress, involved in cancer progression [73]. The excess of reactive oxygen leads to cell damage as well as cancer development [74], the antioxidants often help in protecting against cancer cell formation. On the other hand, GSH is converted to a diverse compound by glutathione-S-transferase, which is responsible in regulating the pathway of mitogen-activated protein (MAP) kinase, responsible for cell survival, as well as being associated with anticancer drug resistance [75]. The probable perturbation of the GSH metabolism by Crotamine opens the possibility of regulating the cellular response. Since, either glutamate or GSH are observed in the NMR spectra, decreased pyroglutamate levels only suggest that Crotamine treatment alters the intracellular redox balance.

Tumor cells can reprogram metabolism in ways that support growth. Following our Crotamine-treated HeLa cells, the BCAAs (Branched-Chain Amino Acid); leucine, isoleucine, and valine, which are essential nutrients for cancer growth, and are present in elevated concentrations impact protein synthesis and degradation [76,77]. Tumor cells uptake BCAA amino acids for protein synthesis or alternatively, oxidizes them to obtain energy [78]. BCAAs are converted into their branched-chain alpha-keto acids by two aminotransferases; one present in the cytosolic space [Branched-chain aminotransferase1 (BCAT1)] and a mitochondrial [branched-chain aminotransferase2 (BCAT2)], the process occurs when the amino group is transferred to the alpha-ketoglutarate, generating glutamate [78,79], affecting the glutathione metabolism, as described earlier. The reduction level of these amino acids in Crotamine treated cells may interfere in the BCAA degradation, and consequently in glutathione metabolism.

In the present study, the lysine concentration decreased under Crotamine treated cells, and we presumed it was being used for cell survival and proliferation. Modifications in lysine are an important functional feature, which regulates cancer development. Dependent of acetyl-CoA, acetylation controlled multiple metabolic processes and lysine acetylation is [80,81] a reversible process, which provide a functional diversity to the protein. When instability in the acetylation is occurring, the stability of the internal environment of the cell can be turned off, as showed in cancer cells [82,83]. We could not test the action of the Crotamine in the acetylation processes; however, lysine is an indispensable amino acid for cell maintenance. The decreased level of lysine in HeLa-treated cell, suggests that this is due to the action of Crotamine in the acetylation process. Histidine is an amino acid that can undergo methylation. 1-methylhistidine, a decreased metabolite in Crotamine treated sample, is a methylated form of histidine, which is frequent in human muscles and urine. 1-methylhistidine is involved in histidine and beta-alanine metabolism [84], the latter, presented altered levels in estrogen receptor for breast cancer subtypes, as well in cancer tissue [84]. Few studies deal with the role of this metabolite in cancer cells.

Formate is a metabolite produce in mammals, being a source of single-carbon group, used for purine synthesis, as well as in methylation [85]. Produced in different tissue from a variety of substrate, it can be synthesized during the catabolism of tryptophan. Formate production is important for a regulatory mechanism, in other words, it is important for the methylation of the DNA, RNA, and proteins [85]. The amount of formate in cancer cells seems to be two times more, compared to normal cells [86,87], and its decrease in the presence of Crotamine, may restrict the supply of the single-carbon group. To conclude, acetate is related to ethanol degradation, and it is further metabolized to acetyl-CoA.

## 5. Conclusions

Following glucose uptake, cancer cell lines preferentially use glycolysis for ATP production, leading to pyruvate and lactate production. Pyruvate is directly linked to the TCA cycle and to valine, leucine and isoleucine metabolism, which in turn, is related to glutathione metabolism and to pyroglutamate levels. Glutathione is the precursor of glycine and thus, glycine, serine, and threonine metabolism are linked to the glutathione and valine, leucine and isoleucine pathways, threonine is metabolized in two ways to be converted in pyruvate. Phenylalanine is the precursor of both pyruvate and tyrosine, which in turn, may be converted to fumarate, a component of the TCA cycle. Finally, aspartate is converted to lysine, threonine and oxaloacetate, also from the TCA cycle.

It is possible to track all observed metabolites following the arrows in figure 7, starting from glucose. Thus, the present results rise the hypothesis that Crotamine may influence the conversion of glucose to pyruvate. Following glucose uptake, it may be consumed by glycolysis, to produce glycogen or may be used in the pentose phosphate pathway [39]. The observation that glucose levels in the medium is reduced in Crotamine-treated sample, indicates that its uptake is increased when compared to untreated HeLa cells. Nevertheless, both pyruvate and lactate, that are synthesized within cell and their surplus are secreted to the medium, are also found to be reduced. This suggests that Crotamine function may be involved in impairing glycolysis. By impairing glycolysis and pyruvate production, it could also bias the TCA cycle, glutathione metabolism and amino acids bio-synthesis and metabolism, as shown in figure 7. However, due to top-down nature of metabolomics approaches, data are inconclusive about how this could be achieved. Crotamine is known to interact with DNA and it could be affecting specific gene expression pathways related to glycolysis or even promoting gene expression in the glycogen synthesis pathway, and in regulating glucose usage in cellular metabolism.

Snake venoms are an interesting source of molecules and are potentially relevant for use in a wide range of biological and clinical applications. Crotamine is a versatile peptide with a variety of biological properties, such as; cell penetration with carrier properties, interaction with DNA and RNA and, as subject of the present study, inhibition of tumor growth and development. The exact mechanism by which Crotamine functions as an anticancer peptide is unclear. In this metabolomics foot printing approach, the levels of certain metabolites were observed to be different following Crotamine treatment of HeLa cells indicating that specific pathways were affected. These results provide an indication of Crotamine function and its mechanism of action in the inhibition of cancer cell growth. Specifically, Crotamine impairs glycolysis, and interferes with other pathways involved glutathione metabolism, the TCA cycle and amino acid biosynthesis and metabolism.

A total of 15 metabolites were changed in the treated sample and provides potential information in cancer progression. Further investigation is needed to validate these initial findings.

## Author Contributions

Conceptualization, M.A.C.; methodology, M.A.C. and F.R.M.; validation, M.A.C., F.R.M., R.J.E., B.S. and M.F.C.; formal analysis, M.A.C., F.R.M., B.S. and M.F.C.; investigation, M.A.C., F.R.M. and B.S.; resources, P.R. and R.K.A.; writing—original draft preparation, M.A.C., F.R.M. and B.S.; writing—review and editing, M.A.C., F.R.M., B.S., M.F.C., R.J.E., P.R. and R.K.A. All authors have read and agreed to the published version of the manuscript.

## Institutional Review Board Statement

Not applicable.

## Informed Consent Statement

Not applicable.

## Data Availability Statement

The data that support the findings of this study are available on request from the corresponding author, [M.A.C].

## Acknowledgments

“We authors would like to dedicate our work to the researchers who gave up their research due the negligence of the Brazilian Federal Government.”

